# Sex-Dependent Vulnerability to PTSD-Like Behaviors in iNOS Knockout Mice

**DOI:** 10.1101/2025.08.27.672399

**Authors:** Bruna F. Ferreira, Isabela Pavan-Silva, Sabrina F. Lisboa

## Abstract

Nitric oxide (NO), mainly produced by neuronal nitric oxide synthase (nNOS) in the brain, has been implicated in stress responses and the pathophysiology of post-traumatic stress disorder (PTSD). Our group previously showed that inducible nitric oxide synthase (iNOS) knockout (KO) male mice exhibit compensatory changes in nNOS expression in the medial prefrontal cortex (mPFC) and impaired fear extinction, suggesting that this genetic model may be relevant for studying PTSD-related phenotypes. Given the frequent comorbidity of PTSD with anxiety and depression, and the marked underrepresentation of females in neuropsychopharmacology research, we performed a behavioral characterization of male and female iNOS KO mice, focusing on aversive memory, anxiety-, and depression-like behaviors. To our knowledge, this is the first systematic behavioral study of female iNOS KO mice, which is particularly relevant given that females are twice as likely to develop psychiatric disorders. We observed that female iNOS KO mice exhibited increased anxiety-like behavior in the elevated plus maze test (EPMT), whereas males showed antidepressant-like behavior in the forced swim test (FST). No general cognitive deficits were found in the Y-maze or object recognition (OR) tests in either sex. However, male iNOS KO mice exhibited deficits in fear extinction memory and extinction retrieval in both contextual and cued fear conditioning. These findings indicate that iNOS KO mice present sex-dependent behavioral phenotypes and may serve as a genetic model to investigate disorders related to fear memory, such as PTSD, and highlight the importance of considering sex as a biological variable in research.

## 1. INTRODUCTION

Post-traumatic stress disorder (PTSD) is associated with fear dysregulation, characterized by exaggerated responses to trauma consolidation and impairments in the extinction of learned aversive responses (Amstadter et al., 2009; Milad et al., 2009). Among the primary comorbidities of PTSD, anxiety disorders and major depressive disorder (MDD) are the most prevalent. Anxiety disorders are characterized by excessive fear and anxiety, while MDD is marked by persistent sadness, loss of interest in previously enjoyable activities, and changes in appetite, sleep, and cognition - symptoms that can significantly reduce quality of life (American Psychiatric Association, 2022). Approximately 50% of individuals diagnosed with PTSD may also meet the criteria for MDD (Rytwinski et al., 2013). Epidemiological studies further indicate that women are twice as likely as men to develop psychiatric disorders such as MDD (Ferrari et al., 2013), panic disorder, and specific phobias (McLean et al., 2011). In the context of PTSD, they are more propense to experience intrusive thoughts, emotional numbing, exaggerated startle responses, and sleep disturbances, whereas men more frequently report symptoms of aggression (Carmassi et al., 2014; Kilpatrick et al., 2013; Perrin et al., 2014; Peters et al., 2006). Despite these well-documented sex differences, a significant underrepresentation of female subjects remains in both clinical and preclinical PTSD research, particularly in those using rodent models (Olff, 2017).

Given that PTSD and its comorbidities are characterized by heightened stress responses (Bin-Jaliah, 2020; Leza et al., 1998; Li et al., 2020), nitric oxide (NO) plays a pivotal role in their pathogenesis by modulating neurotransmitter activity and contributing to oxidative and nitrosative stress (Fronza et al., 2024). At the physiological level, NO regulates the release of neurotransmitters such as glutamate and gamma-aminobutyric acid (GABA) and is involved in key processes including learning, memory formation, sleep, feeding, and pain modulation (Amitai, 2010; Garthwaite, 2019; Prast & Philippu, 2001). This soluble gas is synthesized from the amino acid L-arginine by the enzyme nitric oxide synthase (NOS) in response to calcium (Ca2+) influx or pro-inflammatory cytokines. There are three isoforms of NOS: neuronal (nNOS), expressed in various brain regions; inducible (iNOS), found in macrophages and glial cells; and endothelial (eNOS), predominantly located in endothelial cells (Dhir & Kulkarni, 2011; Esplugues, 2002). Notably, nNOS and iNOS are more closely associated with the nitrergic dysregulation observed in PTSD and its comorbidities (Jankovic et al., 2024).

Particularly, nNOS plays a well-documented role in defensive behaviors and stress-related responses, with its systemic or localized blockade in various brain structures exerting anti-aversive effects and mitigating the consequences of stress exposure (Guimarães et al., 2005; Vila-Verde et al., 2016). Studies have demonstrated that the local injection of a selective nNOS inhibitor into the prelimbic prefrontal cortex (PL PFC) (Moraes Resstel et al., 2008) or dorsal hippocampus (Fabri et al., 2014) of rats reduces fear-related behavior (freezing) in the contextual fear conditioning (CFC) paradigm. Additionally, nNOS knockout (nNOS KO) mice display anxiolytic-like behavior in the elevated plus maze test (EPMT) (Wultsch et al., 2007) and reduced freezing time in CFC (Kelley et al., 2009). Similarly, systemic administration of a preferential nNOS inhibitor in wild-type (WT) animals tested in CFC produces effects comparable to those observed in nNOS KO mice (Kelley et al., 2010). In contrast, iNOS plays a distinct role in stress responses. Its persistent activation, primarily regulated at the transcriptional level and induced by stress exposure, leads to the production of toxic levels of NO in the cerebral cortex and hippocampus (Domingues et al., 2019; Olivenza et al., 2000). This toxicity is mediated by glutamate and other excitatory amino acids, which activate NMDA receptors and trigger the transcription factor NF-kB. Given the distinct but complementary roles of nNOS and iNOS in stress and anxiety-related behaviors, their interplay could provide valuable insights into the mechanisms underlying PTSD maintenance, highlighting potential targets for therapeutic intervention.

Supporting this, iNOS knockout (iNOS KO) animals exhibit reduced brain damage in response to inflammatory stimuli (Kaminska et al., 2016). Moreover, both pharmacological inhibition and genetic deletion of iNOS induce antidepressant-like effects while attenuating the behavioral consequences of stress exposure (Gilhotra & Dhingra, 2009; Montezuma et al., 2012). Additionally, inhibiting iNOS activity in the PL PFC of rats prevents the anxiogenic effects induced by acute stress (Coelho et al., 2022). Thus, while iNOS is traditionally associated with the activation of inflammatory mechanisms (Zlatković & Filipović, 2012), it also plays a role in stress responses. However, seemingly contradictory findings that iNOS KO animals exhibit increased anxiety-like behavior, which becomes even more pronounced one week after stress exposure (Abu-Ghanem et al., 2008; Buskila et al., 2007), as well as heightened freezing behavior in the CFC paradigm (Lisboa et al., 2015). These discrepancies may arise from compensatory adaptations in the nitrergic system within brain structures involved in stress responses. Notably, iNOS KO animals exhibit increased basal activity of the Ca2+-dependent NOS isoforms (nNOS and eNOS) in the PFC and amygdala. The behavioral responses observed in these animals were attenuated by non-selective or preferential nNOS inhibitors, respectively (Buskila et al., 2007; Lisboa et al., 2015). Therefore, iNOS KO animals provide a valuable model for investigating the role of nitrergic signaling and pathways in the persistence of aversive memories, with important implications for PTSD research.

Currently, there is little to no characterization of behavioral responses in female iNOS KO mice. Given the underrepresentation of sex as a biological variable in neuropsychopharmacology research (Cahill, 2012) and the higher prevalence of neuropsychiatric disorders in women (Bao & Swaab, 2011; Merikangas & Almasy, 2020), this study aimed to investigate how biological sex influences the phenotype of iNOS KO animals, particularly in behaviors related to anxiety, depression, fear, and cognitive performance. Furthermore, to the best of our knowledge, this is the first study to compare male and female iNOS KO mice and analyze their behavioral differences in relation to wild-type controls.

## 2. MATERIALS AND METHODS

### 2.1. Animals

This study was performed in 62 male and 68 female iNOS knockout mice (iNOS KO, C57BL/6 background; obtained from the Center for the Development of Special Mice at FMRP/USP – originally acquired from Jackson Laboratory, Bar Harbor, ME, USA), and 59 male and 64 female C57BL/6J control mice (WT), sourced from the Central Animal Facility (FMRP/USP). The animals were housed in ventilated cages with *ad libitum* access to food (Nuvital, Nuvilab, Brazil) and water, maintained on a 12-hour light-dark cycle (lights on at 6:30 a.m.) at a controlled temperature of 24 ± 1°C until they reached the appropriate age for the experiments (8–10 weeks). Animals used in experiments one and three were housed in groups of 6-10 animals per cage (38.9 × 25.1 × 24 cm; Alesco, Brazil), while those in experiment two were housed in groups of 3-5 animals per cage (31.6 × 21.6 × 20.7 cm; Alesco, Brazil).

Procedures were conducted in conformity with the Brazilian Society of Neuroscience and Behavior guidelines for the care and use of laboratory animals, which follow international laws and politics, and were approved by our local ethical committee (nº 20.1.554.60.1 and nº 22.1.757.60.1). All efforts were made to minimize animal suffering and to reduce the number of animals used.

### 2.2. Behavioral Procedures

All behavioral experiments were conducted using male and female WT and iNOS KO mice during the light phase, between 8:00 a.m. and 6:00 p.m.

All apparatuses were cleaned with a 70% ethanol solution between trials to eliminate olfactory clues. Data from the EPMT, OFT, and both contextual and cued fear conditioning paradigms were analyzed offline using AnyMaze software (Stoelting, IL, USA). Data from the FST, ST, Y-maze, and NOR were analyzed manually by a blinded experimenter.

#### 2.2.1. Elevated plus maze test (EPMT)

Anxiety-like behavior was determined using EPMT. EPMT comprised two opposing wooden open arms without side walls (30 x 5 x 15 cm) perpendicular to two closed arms of the same size, all connected by a central platform (5 x 5 cm). The apparatus is elevated 20 cm above the ground. Mice were placed in the center of the maze facing toward one open arm and activity was recorded with an overhead camera for five minutes (Carola et al., 2002). Metrics included the number of entries in the closed arms and the percentages of entries into and time spent in the open arms. The test occurred between 9:00 and 11:00 a.m.

#### 2.2.2. Open field test (OFT)

Locomotor activity and anxiety-like behavior were assessed in OFT, which was conducted in a circular arena enclosed by acrylic walls (30 × 30 cm). Three hours after EPMT, mice were placed in the corner of the arena and allowed to freely explore for six minutes (Becker et al., 2021). Metrics included total distance traveled, distance traveled in the center and periphery, and time spent in the center of the arena. The test occurred between 1:00 and 5:00 p.m.

#### 2.2.3. Forced swimming test (FST)

FST was performed to assess passive behavior (immobility), which is associated with coping strategies in an inescapable situation and can also indicate potential antidepressant effects. Immediately after the OFT, mice were individually placed in an acrylic cylinder (19 × 30 cm) filled with 10 cm of water at 24 ± 1°C for six minutes (Yankelevitch-Yahav et al., 2015). The water was replaced for each animal to maintain a controlled temperature and prevent the influence of alarm substances. The time the animals spent immobile (making only small movements to avoid drowning) was videotaped and manually measured during the first 2-min and last 4-min period. The test occurred between 1:00 and 5:00 p.m.

#### 2.2.4. Splash test (ST)

ST assessed grooming behavior induced by the application of a 10% sucrose solution on the dorsal coat of the animal. The absence of this behavior is associated with anhedonia-like behavior. After applying sucrose solution, mice were placed in acrylic boxes containing 25 g of wood shavings from their home cages (Becker et al., 2021). The latency (time between spray and initiation of grooming) and duration of grooming was recorded for six minutes and manually coded as an index of self-care and motivational behavior. The test occurred between 1:00 and 5:00 p.m.

#### 2.2.5. Contextual Fear Conditioning

The experimental procedure was conducted as depicted in figure 2a. Mice were habituated to a conditioning box (Ugo Basile Fear conditioning system) and subsequently received three inescapable footshocks (0.75 mA; 2s each) through a metal grid (day 1) to assess the behavioral fear response (freezing) (Lisboa et al., 2018). On the following day (day 2), mice were re-exposed to the box for assessment of extinction acquisition for 20 min. After 24h (day 3), the extinction memory retrieval was assessed for 5 min in the same chamber. All tests occurred between 9:00 and 11:00 a.m.

#### 2.2.6. Cued Fear Conditioning

The experimental procedure was conducted as illustrated in figure 2e. On day 1, mice underwent six conditioning trials in context A. Each trial consisted of a 30-second presentation of the conditioned stimulus (CS; 70 dB, 1 Hz tone), which co-terminated with a 1-second unconditioned stimulus (US; 0.45 mA footshock). After 24h (day 2), mice underwent extinction training in context B, which included olfactory enrichment (10% vanilla), consisting of 18 CS presentations. On day 3, an extinction retrieval session was performed, involving 9 CS presentations in context B (Huang et al., 2020). All tests occurred between 9:00 and 11:00 a.m.

#### 2.2.7. Y-maze spontaneous alternation test

The Y-maze test takes advantage of the innate investigative nature of rodents to explore new environments to assess short-term spatial working memory. The apparatus consists of three identical Plexiglas arms (36 x 6 x 16 cm) separated by a 120º angle. Each animal was placed at the end of one arm and allowed to move freely through the arms during an eight-minute session (Kraeuter et al., 2019). All tests occurred between 9:00 and 11:00 a.m. Successful arm alternation was defined as the animals entering different arms successively (e.g., ABC, CBA, ACB), counted from the first arm entry and always in sequences of three. An entry and an exit of an arm were defined as when more than 50% of the mice body crossed the boundary of the center. The percentage of spontaneous alternation was calculated as follows:

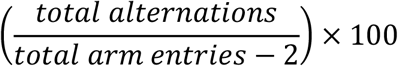

#### 2.2.8 Novel object recognition (NOR)

NOR is used to evaluate episodic declarative memory in rodents. This test relies on the animal’s natural tendency to explore a novel object more than a familiar one in a known context. This test consisted of a habituation phase, a training phase, a short-term testing phase, and a long-term testing phase. The animals were habituated to a square acrylic arena (40 x 30 cm) for 30 minutes. On the following day, the animals were re-exposed to the box with two identical objects (A+A) for free exploration over five minutes (acquisition session). After 90 minutes, the short-term memory (STM) test session was conducted, during which one of the objects was replaced (A+B) and allowed to be explored for five minutes. After 24 hours, the long-term memory (LTM) test session was performed. For the LTM session, object A was retained, and object B was replaced with a new object (A+C) (Moore et al., 2013; Wang & Han, 2018). Animals that did not interact with the objects for at least five seconds were excluded from the test. The time spent exploring each object was recorded and expressed as a recognition index (RI), where RI = time exploring novel object/(time exploring novel object + time exploring familiar object)x100. Habituation phase occurred between 1:00 and 5:00 p.m.

### 2.3. Statistical analyses

Data were initially tested for normality using the Shapiro-Wilk test and for homogeneity of variances using the Brown-Forsythe test. Normally distributed and homogenous data were analyzed using two-way ANOVA or Student’s t-test. Normally distributed but non-homogenous data were analyzed using one-way ANOVA with Brown-Forsythe correction or Welch’s adjustment. Data that did not follow a normal distribution were analyzed using the non-parametric Kruskal-Wallis test or the Mann-Whitney test. For temporal analyses, data were first tested for sphericity using Mauchly’s test. If sphericity was violated, the Geisser-Greenhouse correction was applied, followed by repeated-measures ANOVA.

When repeated-measures ANOVA or two-way ANOVA identified an interaction effect between factors, data were subjected to multiple comparisons using Tukey’s or Dunnett’s post-hoc tests to evaluate intergroup and intragroup effects. For one-way ANOVA or Kruskal-Wallis tests that identified group differences, data were compared using Dunnett’s or Dunn’s post-hoc tests, respectively. The significance level was set at p ≤ 0.05.

Analyses were performed using GraphPad Prism software, version 10.0 (GraphPad Software; San Diego, CA, USA). Results are presented as mean ± standard error of the mean (SEM). The symbols used were: @ for differences related to sex (male and female); # for differences related to genotype (WT and iNOS KO); and * for differences related to time.

## 3. RESULTS

### 3.1. Experiment 1 – Evaluation of behavioral responses in male and female WT and iNOS KO mice subjected to anxiety and depression tests

To evaluate whether biological sex influences the potential anxiety-like and antidepressant-like phenotype of iNOS KO mice, we conducted a series of behavioral tests for anxiety-like and depressive-like behaviors in the following order: elevated-plus maze test (EPMT), open-field test (OFT), and forced swimming test (FST). Additionally, an independent group of mice was subjected exclusively to the splash test (ST).

In the EPM, female iNOS KO mice showed a reduced percentage of entries into the open arms (figure 1c; two-way ANOVA – sex: F_(1,49)_ = 14.72, p = 0.0004; genotype: F_(1,49)_ = 6.58, p = 0.01; interaction: F_(1,49)_ = 4.43, p = 0.04) compared to male iNOS KO mice (p = 0.0009; Tukey’s post-hoc test) and female WT mice (p = 0.007; Tukey’s post-hoc test). The analysis of the percentage of time spent in the open arms (figure 1b) revealed significant effects of sex (F_(1,49)_ = 7.89, p = 0.007) and genotype (F_(1,49)_ = 13.85, p = 0.0005) but no interaction between factors (F_(1,49)_ = 0.007, p = 0.93), according to two-way ANOVA. Overall, females spent less time in the open arms than males, and this effect was observed across iNOS KO and WT groups. Additionally, male iNOS KO mice exhibited fewer entries into the closed arms (Figure 1d; Brown-Forsythe ANOVA – F_(3,30)_ = 8.17, p = 0.0004) compared to WT males (p < 0.0001; Dunnett’s post-hoc test), while female iNOS KO mice showed a higher number of entries compared to male iNOS KO mice (p = 0.03; Dunnett’s post-hoc test).

**Figure 1.**
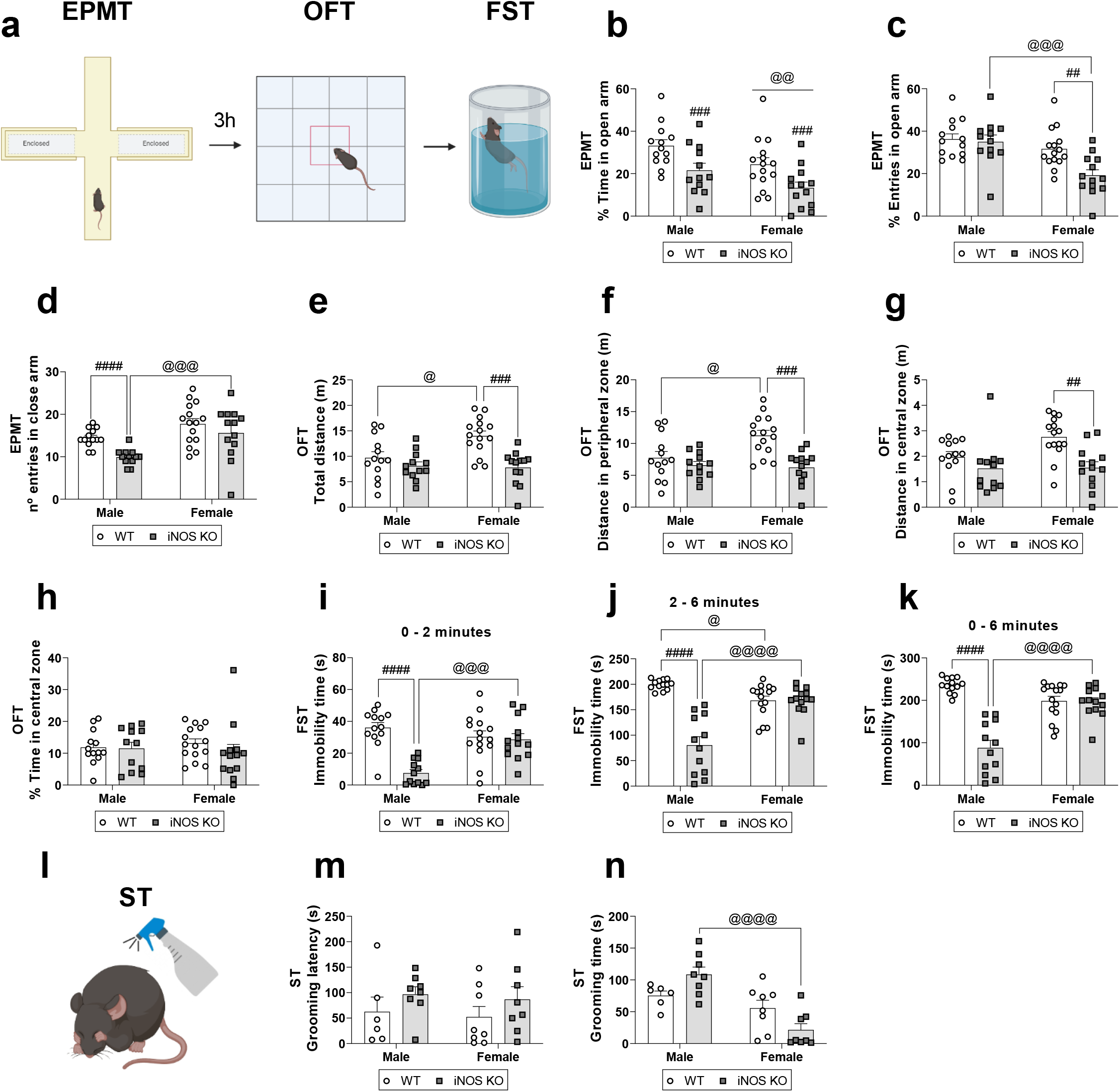
Comparison of anxiety- and antidepressant-like behaviors between male and female WT and iNOS KO mice. (a) Graphical representation of the experimental design. (b) Percentage of time spent in the open arms of the EPM. (c) Percentage of entries into the open arms of the EPM. (d) Number of entries into the closed arms of the EPM. (e) Total distance traveled in the OFT. (f) Distance traveled in the periphery of the OFT. (g) Distance traveled in the center of the OFT. (h) Time spent in the center of the OFT. (i) Immobility time during the first two minutes of the FST. (j) Immobility time during the last four minutes of the FST. (k) Total immobility time in the FST. (l) Graphical representation of ST. (m) Latency to grooming in the ST. (n) Time spent grooming in the ST. Data are presented as Mean ± SEM and analyzed using two-way ANOVA (A, B, D, and E), one-way ANOVA with Brown-Forsythe correction (C), or Kruskal-Wallis nonparametric test (F, G, H, I, J, K, and L). A total of 8–15 animals per group were used. @p < 0.05, @@p < 0.01, @@@p < 0.001, or @@@@p < 0.0001 compared between sexes; ##p < 0.01, ###p < 0.001, or ####p < 0.0001 compared between genotypes. EPMT: Elevated Plus Maze Test; OFT: Open Field Test; FST: Forced Swim Test; ST: Splash Test.

In the OFT, female WT mice exhibited hyperlocomotion when compared to male WT mice (p = 0.01; Tukey’s post-hoc test), as indicated by increased total distance traveled (figure 1e; two-way ANOVA – sex: F_(1,49)_ = 4.15, p = 0.047; genotype: F_(1,49)_ = 16.09, p = 0.0002; interaction: F_(1,49)_ = 5.61, p = 0.02). This effect was also noted in the periphery of the apparatus (figure 1f; two-way ANOVA – sex: F_(1,49)_ = 3.85, p = 0.055; genotype: F_(1,49)_ = 14.77, p = 0.0003; interaction: F_(1,49)_ = 5.77, p = 0.02). In contrast, female iNOS KO mice displayed hypolocomotion compared to WT females (p = 0.0002; Tukey’s post-hoc test) for these same parameters. Hypolocomotion was also evident in the distance traveled in the center of the apparatus (figure 1g; Kruskal-Wallis non-parametric test – K = 16.53, p = 0.0009), with a significant reduction compared to WT females (p = 0.004; Dunn’s post-hoc test). However, no significant differences were observed for the time spent in the center (Figure 1h; Kruskal-Wallis non-parametric test – K = 3.9, p = 0.27).

In the FST, male iNOS KO mice showed reduced immobility time during the first two minutes (figure 1i; Kruskal-Wallis non-parametric test – K = 23.26; p < 0.0001), the last four minutes (figure 1j; Kruskal-Wallis non-parametric test – K = 32.32; p < 0.0001), and the total test duration (figure 1k; Kruskal-Wallis non-parametric test – K = 30.7, p < 0.0001), compared to female iNOS KO mice (p = 0.006, p = 0.02, and p = 0.008, respectively; Dunn’s post-hoc test) and male WT mice (p < 0.0001; Dunn’s post-hoc test). A reduction in immobility time was also observed in female WT mice compared to males during the last four minutes of the test (figure 1j; p = 0.03; Dunn’s post-hoc test).

Finally, in the ST, female iNOS KO mice spent less time in grooming (figure 1m; Kruskal-Wallis non-parametric test – K = 16.61, p = 0.0009) compared to male iNOS KO mice (p = 0.0009; Dunn’s post-hoc test). However, no differences were observed in the latency to initiate grooming between groups (figure 1n; Kruskal-Wallis non-parametric test – K = 6.8, p = 0.08).

### 3.2. Experiment 2 – Evaluation of contextual and cued fear conditioning responses in male and female WT and iNOS KO mice

In the second experimental protocol, mice were exposed to an aversive stimulus (foot shock), and their fear response was evaluated. This protocol involved two distinct groups: Group A, consisted of mice exposed to the contextual fear conditioning paradigm, whereas Group B consisted of mice subjected to the cued fear conditioning paradigm. This approach was designed to compare the effects of the two paradigms on the fear response of iNOS KO mice.

#### 3.2.1. Contextual fear conditioning: acquisition, extinction, and retrieval

During the conditioning session of contextual fear conditioning (day 1; figure 2b), repeated-measures ANOVA revealed a significant effect of time (F_(2.55,112.07)_ = 127.3, p < 0.001), genotype (F_(1,44)_ = 5.2, p = 0.03), and a time × genotype interaction (F_(2.55,112.07)_ = 3.18, p = 0.03). No significant effects were observed for sex (F_(1,44)_ = 1.53, p = 0.23), sex × genotype interaction (F_(1,44)_ = 0.001, p = 0.98), time × sex interaction (F_(2.55,112.07)_ = 1.58, p = 0.20), or the interaction between all three factors (F_(2.55,112.07)_ = 0.65, p = 0.56). These findings indicate that iNOS KO mice exhibited a higher percentage of freezing compared to WT mice, although all groups successfully acquired fear memory.

**Figure 2.**
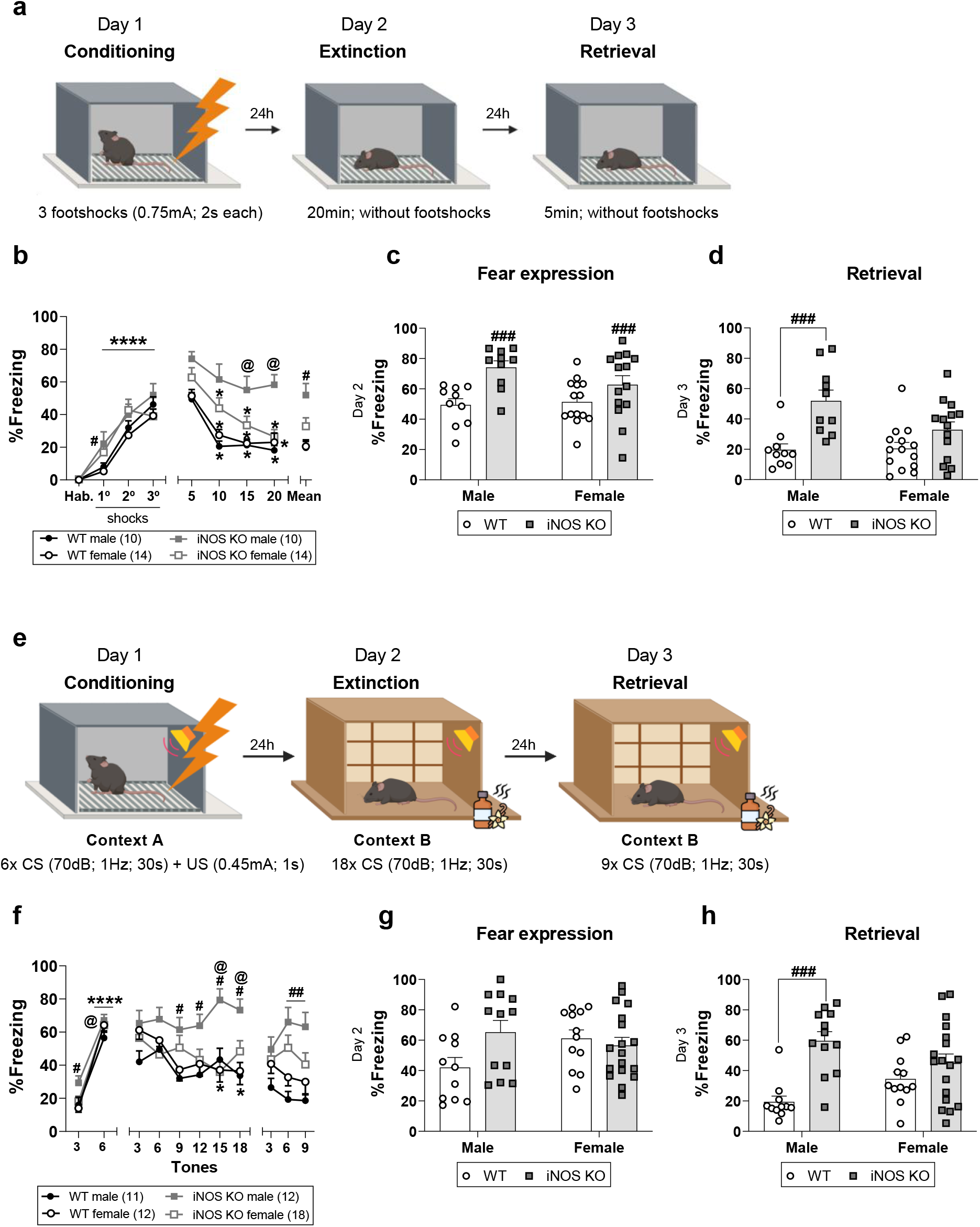
Comparison of contextual and cued fear conditioning responses between male and female WT and iNOS KO mice. (a) Graphical representation of contextual fear conditioning paradigm. (b) Conditioning and extinction over time in contextual fear conditioning. (c) Fear expression in contextual fear conditioning. (d) Extinction retrieval in contextual fear conditioning. (e) Graphical representation of cued fear conditioning paradigm. (f) Conditioning, extinction, and retrieval over time in cued fear conditioning. (g) Fear expression in cued fear conditioning. (h) Total extinction retrieval in cued fear conditioning. Data are presented as Mean ± SEM and were analyzed using repeated-measures ANOVA (b and f), two-way ANOVA (c), or the non-parametric Kruskal-Wallis test (d, g, and h). A total of 10–18 animals per group were used. @p < 0.05 compared between sexes; #p < 0.05, ##p < 0.01, ###p < 0.001 compared between genotypes. *p < 0.05, ****p < 0.0001 for time comparisons.

On day 2, both male and female iNOS KO mice demonstrated higher fear expression compared to WT mice (Figure 2c; two-way ANOVA – sex: F_(1,44)_ = 0.93, p = 0.34; genotype: F_(1,44)_ = 13.6, p = 0.0006; sex × genotype interaction: F_(1,44)_ = 1.9, p = 0.18). During the extinction session, there was a reduction in freezing percentage in WT mice and female iNOS KO mice, but not in male iNOS KO mice (day 2; Figure 2b; repeated-measures ANOVA – time: F(2.56,112.71) = 42.42, p < 0.001; genotype: F(1,44) = 31.12, p < 0.001; sex: F(1,44) = 4.31, p = 0.04; sex × genotype interaction: F(1,44) = 8.95, p = 0.005). There were no significant effects for time × sex interaction (F(2.56,112.71) = 1.09, p = 0.35), time × genotype interaction (F(2.56,112.71) = 1.24, p = 0.30), or the interaction between time, sex, and genotype (F(2.56,112.71) = 1.45, p = 0.24).

During the recall session (day 3; figure 2d), male iNOS KO mice exhibited a higher percentage of freezing compared to male WT mice (non-parametric Kruskal-Wallis test – K = 14.85, p = 0.002; Dunn’s post hoc test – WT vs. iNOS KO males: p = 0.005). These results suggest that WT mice and female iNOS KO mice successfully acquired and recalled extinction memory, while male iNOS KO mice exhibited deficits in these processes.

#### 3.2.2. Cued fear conditioning: acquisition, extinction, and retrieval

During the conditioning phase of cued fear conditioning (day 1; figure 2f), all groups acquired fear memory, as evidenced by increased freezing levels throughout the session. However, this increase was more pronounced in iNOS KO mice (repeated-measures ANOVA – time: F(_1,49)_ = 682.41, p < 0.001; genotype: F(_1,49)_ = 4.31, p = 0.04; sex: F_(1,49)_ = 0.38, p = 0.54). No significant interactions were observed between sex × genotype (F_(1,49)_ = 2.22, p = 0.14), time × genotype (F_(1,49)_ = 1.07, p = 0.31), or time × sex × genotype (F_(1,49)_ = 0.03, p = 0.86), though there was a significant interaction between time and sex (F_(1,49)_ = 7.36, p = 0.009).

On day 2, there wasn’t alterations in fear expression across groups, with no significant differences in freezing percentages (figure 2g; Kruskal-Wallis test – K = 5.59, p = 0.13). However, during the extinction phase (day 2; figure 2f), repeated-measures ANOVA revealed a significant effect of genotype (F_(1,49)_ = 9.89, p = 0.003) and a trend for an effect of time (F_(5,45)_ = 2.05, p = 0.09). No main effect of sex was detected (F_(1,49)_ = 2.58, p = 0.12). Interactions were observed between sex and genotype (F_(1,49)_ = 7.32, p = 0.009) and between time and sex (F_(5,45)_ = 3.53, p = 0.009), but not between time and genotype (F(_5,45)_ = 1.54, p = 0.20) or among all three factors (F_(5,45)_ = 0.36, p = 0.87). These findings indicate that male iNOS KO mice exhibited higher freezing levels compared to male WT and female iNOS KO mice.

Extinction memory retrieval was analyzed in two ways. In the overall analysis (figure 2h), male iNOS KO mice displayed significantly higher freezing percentages than male WT mice (Kruskal-Wallis test – K = 16.96, p = 0.0007; Dunn’s post hoc test – WT vs. iNOS KO males: p = 0.0002). When time was considered (day 3; figure 2f), repeated-measures ANOVA showed a significant effect of genotype (F(_1,49)_ = 18.04, p < 0.001), while no effects were found for sex (F_(1,49)_ = 0.02, p = 0.88) or time (F_(2,48)_ = 0.56, p = 0.57). There were interactions between sex and genotype (F_(1,49)_ = 5.93, p = 0.02) and between time and genotype (F_(2,48)_ = 5.34, p = 0.008), but not between time and sex (F_(2,48)_ = 0.84, p = 0.47) or among all three factors (F(_2,48)_ = 0.46, p = 0.63).

The results also highlight that male iNOS KO mice maintained higher freezing levels compared to other groups throughout the session. These findings suggest deficits in both acquisition of fear extinction and retrieval of extinction memory in male iNOS KO mice, mirroring the effects observed in contextual fear conditioning.

### 3.3. Experiment 3 – Evaluation of cognitive performance in male and female WT and iNOS KO mice across non-aversive memory tasks

Given the extinction deficits of fear memory observed in male iNOS KO mice, the next step was to assess other forms of memory to determine whether these animals exhibit generalized cognitive impairment or specific deficits tied to aversive experiences. In this context, the third experiment evaluated spatial working memory using the Y-maze test and episodic working memory, as well as short- and long-term memory, using NOR.

In the Y-maze test (figure 3b), females exhibited a lower percentage of spontaneous alternations compared to males, regardless of genotype (two-way ANOVA – sex: F_(1,52)_ = 9.26, p = 0.004; genotype: F_(1,52)_ = 0.41, p = 0.53; interaction: F_(1,52)_ = 0.34, p = 0.56). These results suggest that iNOS KO mice did not display deficits in spatial working memory formation, although females performed differently than males.

**Figure 3.**
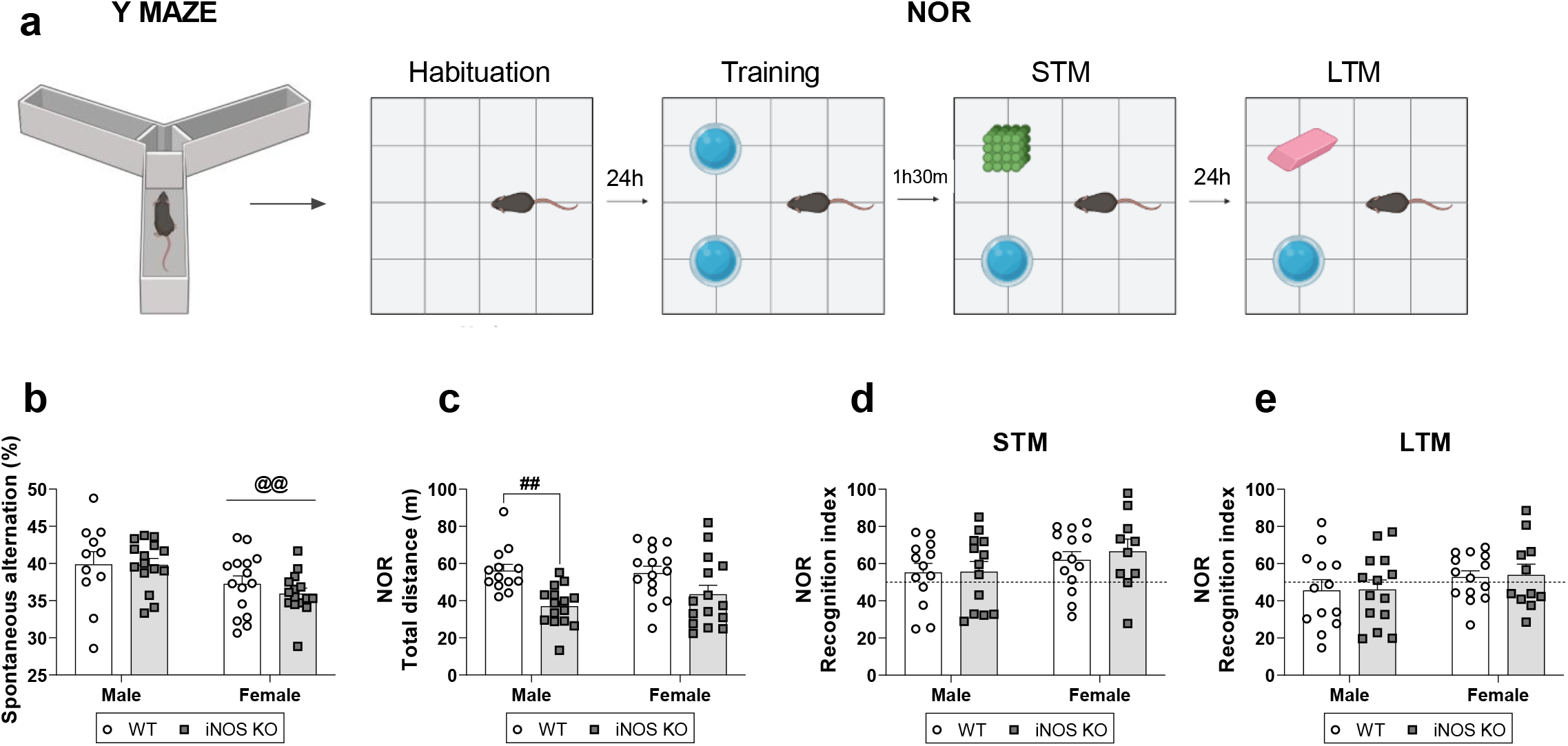
Comparison of cognitive performance between male and female WT and iNOS KO mice. (a) Graphical representation of the experimental design. (b) Percentage of spontaneous alternation of the Y-maze test. (c) Total distance traveled in the habituation session of the NOR test. (d) Recognition index in the STM session of the NOR test. (e) Recognition index in the LTM session of the NOR test. Data are presented as Mean ± SEM and analyzed using two-way ANOVA (b, d, and e) or Kruskal-Wallis nonparametric test (c). A total of 11–15 animals per group were used. @@p < 0.01 compared between sexes; ##p < 0.01 compared between genotypes. NOR: Novel Object Recognition; STM: Short-Term Memory; LTM: Long-Term Memory.

Following Y-maze, mice underwent the NOR test. During the habituation session (figure 3c), iNOS KO male mice demonstrated hypolocomotion compared to WT male mice, as indicated by reduced total distance traveled (non-parametric Kruskal-Wallis test – K = 15.66; p = 0.001; Dunn’s post hoc – WT vs. iNOS KO male: p = 0.006). In the STM session (figure 3d), there were no differences in the recognition index among the groups (two-way ANOVA – sex: F_(1,46)_ = 2.83, p = 0.1; genotype: F_(1,46)_ = 0.22, p = 0.64; interaction: F_(1,46)_ = 0.16, p = 0.69). Similarly, during the LTM evaluation (figure 3e), the recognition index showed no significant differences between groups (two-way ANOVA – sex: F_(1,48)_ = 2.28, p = 0.14; genotype: F_(1,48)_ = 0.03, p = 0.87; interaction: F_(1,48)_ = 0.005, p = 0.95).

These findings indicate that the cognitive performance of iNOS KO mice remains unimpaired, even though females exhibit different behavioral patterns compared to males.

## 4. DISCUSSION

In this study, we compared the behavioral phenotype of male and female WT and iNOS KO mice, as such a comparison has not yet been described in literature. Previous evidence suggests that male iNOS KO mice exhibit impaired fear memory extinction (Lisboa et al., 2015), anxiety-like behavior (Abu-Ghanem et al., 2008; Buskila et al., 2007; Fernandes et al., 2021), and, controversially, antidepressant-like behavior (Fernandes et al., 2021; Issy et al., 2018; Montezuma et al., 2012). However, these behaviors have not been previously analyzed in female mice. To the best of our knowledge, this is the first study to characterize the behavior of female iNOS KO mice in tests evaluating anxiety- and depressive-like phenotypes, as well as cognitive performance. This is particularly relevant for understanding stress-related disorders, such as PTSD, which is twice as common in women (Kilpatrick et al., 2013; Perrin et al., 2014).

We observed that female iNOS KO mice exhibited anxiety-like behavior compared to both male iNOS KO and female WT mice. This was evidenced by a 45% reduction in time spent and fewer entries into the open arms of the EPMT. Traditionally, the estrous cycle has been considered a major source of increased variability in female subjects, potentially influencing animal behavior (Chari et al., 2020; Tropp & Markus, 2001; Walf et al., 2009). As a result, it has been widely believed that females exhibit higher behavioral fluctuations than males. However, recent studies challenge this notion, suggesting that female mice display more homogeneous behavior than males (Kaluve et al., 2022; Levy et al., 2023), and that female sex or endogenous estradiol levels do not have a significant impact on behavioral variability (Beery, 2018; Tsao et al., 2023). Therefore, the differences in anxiety-like behaviors observed in this study could be attributed to other factors, including the genetic background, environmental conditions, and handling procedures.

Female iNOS KO mice have been shown to exhibit hypolocomotion (Casarotto et al., 2018). Corroborating with these findings, we observed that female iNOS KO mice also displayed reduced locomotion compared to female WT mice in the OFT. Additionally, female WT mice exhibited a 44% increase in locomotor activity compared to male WT mice. This difference may be attributed to the tendency of female mice to exhibit hyperlocomotion under specific conditions, such as exposure to novel environments, stress protocols, or as a result of genetic modifications (Bishnoi et al., 2021; Caldarone et al., 2008; Tucker et al., 2016). On the other hand, male iNOS KO mice spent less time in the open arms of the EPMT, although there was no significant difference in the percentage of entries into the open arms. They also showed a reduced number of entries into the closed arms, suggesting hypolocomotion. This locomotor impairment has been previously reported in other studies (Fernandes et al., 2021; Issy et al., 2018). However, in the OFT, male iNOS KO mice did not exhibit differences in total distance traveled, nor in the distance covered in the peripheral and central zones. Therefore, consistent with previous studies (Abu-Ghanem et al., 2008; Fernandes et al., 2021), we conclude that these mice also display anxiety-like behavior.

Considering depressive-like behaviors, we observed that female iNOS KO mice spent 76% less time grooming in the ST compared to male iNOS KO mice, suggesting a depressive-like phenotype. However, this pattern was not observed in the FST. Additionally, in the FST, female WT mice exhibited a 15% reduction in immobility time compared to male WT mice. Although both FST and ST are behavioral assays used to assess depressive-like behaviors, they differ significantly in their underlying principles (Gencturk & Unal, 2024). While the FST evaluates passive coping strategies (immobility), the ST assesses active coping strategies (grooming and exploration). Thus, it is possible that female iNOS KO are more sensitive to deficits in active coping mechanisms rather than passive coping strategies. Meanwhile, male iNOS KO mice displayed an antidepressant-like phenotype, as evidenced by a more than 50% decrease in immobility time throughout the FST session. These findings are consistent with previous literature, which reports that both genetic deletion and pharmacological inhibition of iNOS produce similar effects in the FST (Montezuma et al., 2012). Interestingly, studies have also shown that increased iNOS expression in the hippocampus is associated with depressive-like behaviors in the FST and the tail suspension test (TST) (de Sá-Calçada et al., 2015). This raises the possibility that the absence of iNOS may confer a protective effect against depressive-like behaviors, as suggested by the reduced immobility observed in our study.

This study also demonstrated that only male iNOS KO mice exhibited deficits in the acquisition and retrieval of extinction in both contextual and cued fear conditioning paradigms. These deficits were evidenced by an increased freezing percentage compared to male WT mice. Meanwhile, female iNOS KO mice successfully exhibited acquisition and retrieval of extinction in both paradigms tested. The literature is conflicting regarding which sex is more prone to exhibit higher levels of fear, with some studies reporting greater fear in males (Barker & Galea, 2010; Clark et al., 2019; Daviu et al., 2014) and others in females (Fenton et al., 2014, 2016). These discrepancies likely reflect the influence of multiple factors. For instance, evidence indicates that the behavioral expression of fear differs between sexes (Borkar et al., 2020; Gruene et al., 2015; Russo & Parsons, 2021). This implies that, in some cases, observed sex differences in fear conditioning may be attributable to variations in behavioral performance rather than differences in learning itself.

Using the same conditioning protocol applied in this study, previous research has shown that male iNOS KO mice display extinction deficits in the contextual fear conditioning paradigm (Lisboa et al., 2015). While we conducted a single 20-minute extinction session, Lisboa et al. employed four extinction sessions of five minutes each, conducted 24, 48, 72, and 96 hours after the conditioning session. Notably, they observed that the extinction deficits in male iNOS KO mice were exacerbated across all sessions. Taken together, these findings reinforce that male iNOS KO mice exhibit deficits in the extinction of fear memory and that these deficits are robust and persist regardless of variations in extinction training.

Individuals with PTSD often experience disturbances in memory function (Petzold & Bunzeck, 2022; Scott et al., 2015). Memory is a complex cognitive function that enables individuals to encode, store, and retrieve information (Zlotnik & Vansintjan, 2019). It is a dynamic process that evolves over time based on new experiences and can be disrupted in conditions such as PTSD (Cobb Scott et al., 2016; Vasterling et al., 2002) and depression (American Psychiatric Association, 2022; Rhee et al., 2024). Memory is typically categorized into three main types: sensory memory, working memory and long-term memory. Long-term memory, in turn, is subdivided into declarative episodic memory and non-declarative associative memory (Camina & Güell, 2017).

Given that male iNOS KO mice exhibited extinction deficits in fear memory, a form of associative learning, we investigated whether other cognitive domains were also compromised, which could indicate a generalized dysfunction. However, spatial working memory, declarative episodic memory, as well as their discriminatory ability were not affected. Since cognitive performance, assessed in Y-maze and NOR test, was preserved in male or female iNOS KO mice, our results suggest that the memory deficits observed in male iNOS KO mice may be restricted to fear memory, at least within the evaluated time frame. It is important to consider that different types of memory can be modulated by distinct NO-dependent mechanisms. Corroborating our findings, previous studies indicate that NO plays a more critical role in emotional memory processes, such as fear conditioning, compared to spatial working memory or object recognition (Da Cunha et al., 2005; Oosthuizen et al., 2005; Schafe et al., 2005). Our results align with the literature, as male iNOS KO mice did not exhibit deficits in spatial memory in Morris water maze (Medeiros et al., 2007) or in declarative memory in the NOR test (Abu-Ghanem et al., 2008). Additionally, male iNOS KO mice only displayed motor and spatial working memory impairments following a traumatic brain injury (Sinz et al., 1999). Given that individuals with PTSD can present cognitive deficits (Prieto et al., 2023; Roberts et al., 2022), a later assessment of cognitive function in these animals might reveal such alterations - an aspect that has yet to be investigated.

## 5. CONCLUSION

Our findings expand the knowledge of the distinct behavioral phenotypes of male and female iNOS KO mice, particularly regarding anxiety, depression, fear, and cognitive performance. While female iNOS KO mice exhibited increased anxiety-like behaviors, male iNOS KO mice showed impairments in the acquisition and retrieval of extinction memory. These findings align with previous studies highlighting the role of NO in emotional memory. However, the observed deficits appear to be specific to fear memory, as neither sex exhibited alterations in general cognitive performance. Overall, our findings reinforce the existence of behavioral differences between males and females, which may be influenced by brain sexual dimorphism, a hypothesis that remains to be investigated. This underscores the importance of including both sexes in behavioral studies to ensure a comprehensive understanding of neurobiological mechanisms. Furthermore, given the alterations observed in male iNOS KO mice, this study supports the potential use of these animals as a model for investigating disorders such as PTSD.

## Acknowledgements

We thank Flavia C. Salatta and Miriam Contim Melo for technical assistance. We also acknowledge the use of AI language model ChatGPT (OpenAI) for assistance with grammar and style revision. The authors are fully responsible for the content and interpretations presented.

## Authors’ contributions

BFF and SL designed the study. BFF and IPS performed the experiments. BFF performed the data analysis. BFF, IPS and SFL wrote the manuscript. SFL supervised the project. All authors revised and have approved the article.

## Ethical standard

The authors assert that all procedures performed in this work comply with the ethical standards of the relevant national and institutional committees on Animal Use Ethics. The work was approved by the Animal Use Ethics Commission (CEUA) of School of Pharmaceutical Sciences of Ribeirao Preto (Approval nº: 20.1.554.60.1 and 22.1.757.60.1).

